# A Tug of War Between Condensate Phases in a Minimal Macromolecular System

**DOI:** 10.1101/2020.01.29.925099

**Authors:** Archishman Ghosh, Xiaojia Zhang, Huan-Xiang Zhou

**Affiliations:** Department of Chemistry, University of Illinois at Chicago, Chicago, IL 60607; Department of Physics, University of Illinois at Chicago, Chicago, IL 60607

## Abstract

Membraneless organelles formed via liquid-liquid phase separation (LLPS) contain a multitude of macromolecular species. A few of these species drive LLPS while most serve as regulators. The LLPS of SH3_5_ (S) and PRM_5_ (P), two oppositely charged protein constructs, was promoted by a polyanion heparin (H) but suppressed by a cationic protein lysozyme (L). Here, using these four components alone, we demonstrate complex phase behaviors associated with membraneless organelles and uncover the underlying physical rules. The S:P, S:L, and P:H binaries form droplets, but the H:L binary forms precipitates, therefore setting off a tug of water between different phases within the S:P:H:L quaternary. We observe dissolution of precipitates upon compositional change, transformation from precipitates to droplet-like condensates over time, and segregation of S:L-rich and P:H-rich foci inside droplet-like condensates. A minimal macromolecular system can thus recapitulate membraneless organelles in essential ways and provide crucial physical understanding.

## Introduction

Membraneless organelles, formed via liquid-liquid phase separation^1^, mediate a host of cellular functions ranging from stress response to ribosome biogenesis, but are also prone to aberrant transitions leading to diseases^2–4^. They assembly and disassemble in response to environmental changes and during cell cycles and development. Typically comprising dozens to hundreds of macromolecular components^5–9^, with many more in the surrounding solution and ready to be recruited, these bimolecular condensates are far from the simple picture of a homogeneous mixture. Indeed, multiple phases with distinct material properties coexist in stress granules^8^, nucleoli^10^, paraspeckles^11^, nuclear speckles^12^, and P granules^13^. Moreover, the material states of many membraneless organelles evolve over time, some for normal functions and others as aberrant transitions^14^. With different phases occurring in space and over time, along with many triggers (e.g., expression levels, posttranslational modifications, disease-associated mutations), there are constant battles between phases. Here we used a four-component macromolecular system to reconstitute some of these complex behaviors and uncover the underlying physical rules.

Droplets are a common, liquid phase of bimolecular condensates, and can assemble or disassemble by a simple compositional change, i.e., when the components go above or below their threshold concentrations^15, 16^. When a macromolecular mixture can exist in multiple condensate phases, each favored by a different composition, a compositional change can likewise result in the changeover from one condensate phase to another. For example, the binary mixtures of nuleophosmin and fibrillarin, two protein components of nucleoli, can form fibrillarin-rich droplets, or nuleophosmin-rich droplets, or droplets within droplets (where fibrillarin droplets are immiscible with but embedded inside nuleophosmin droplets), depending on the concentrations of the two components^10^. Similarly, the binary mixtures of speckle-type POZ protein (SPOP) and the death-domain-associated protein (DAXX) form network-like precipitates at low DAXX concentrations but droplets at high DAXX concentrations^17^. This changeover happens when the condensation shifts from being driven by stronger SPOP-DAXX interactions to by weaker DAXX-DAXX interactions. The role of intermolecular interaction strength in the competition between phases is further supported by observations that condensates formed by a protein:DNA binary change from droplets into network-like precipitates when electrostatic repulsion is reduced by mutations, and the precipitates revert back to droplets when the electrostatic attraction is weakened by increasing the salt concentration^18^. A protein-arginine repeat peptide forms droplets when mixed with any one of three homopolymeric RNAs (poly-rA, rU, rC) but forms network-like precipitates when mixed with the fourth, poly-rG^19^. The explanation for the exception was that poly-rG forms quadruplexes, suggesting that the degree of structural compactness is also a determinant for condensate phases.

After phase separation, droplets formed by several component ribonucleoproteins (FUS, hnRNPA1, hnRNPA2, TDP43, and TIA1) of stress granules undergo a further transition into a solid-like state, such as irreversible gels or fibrils^20–30^. This process is undoubtedly facilitated by the high protein concentrations inside droplets and, strikingly, is promoted or accelerated by mutations associated with neurodegenerative diseases (including amyotrophic lateral sclerosis (ALS) and frontotemporal dementia). Similar observations have been made on tau, a protein associated with Alzheimer’s disease, again suggesting that droplets can serve as an intermediate toward protein aggregation^31^. ALS-associated, misfolded SOD1 mutants accumulate in stress granules, where they aggregate, distribute inhomogeneously, and displace component proteins, thereby making the stress granules more solid-like^32^.

The liquid-to-solid transition of biomolecular condensates poses a challenge for their disassembly and hence may lead to the accumulation of pathological inclusions. Several studies have demonstrated the roles of disaggregases. For example, chaperones such as HSP70 are recruited into stress granules containing misfolded proteins to prevent their accumulation and thereby promote the disassembly of these stress granules^32^. Likewise, while acidic pH-induced Pub1 condensates are reversible gels that readily dissolve upon returning to neutral pH, heat-induced Pub1 condensates are solid-like and their dissolution requires the chaperone HSP104^33^. Maharana et al.^34^ proposed that high levels of RNA, in essence acting as a disaggregase, keep ribonucleoproteins soluble in the nucleus. Supporting this proposition, digestion of RNA by microinjection of RNase A into the nucleus triggers the phase separation of FUS and other ribonucleoproteins; conversely, mutations impairing RNA binding result in solid-like FUS inclusions in the cytoplasm. Similarly, mutations impairing RNA binding result in solid-like TDP43 inclusions in both the cytoplasm and the nucleus, and an oligonucleotide that specifically binds TDP43 can rescue aberrant phase transitions^35^. How disaggregases work is largely unknown. Of the two nucleolar components, nuleophosmin droplets remain liquid-like over time but fibrillarin droplets become more solid-like^10^. It has been suggested that, inside cells, an unknown, ATP-dependent process actively maintains the internal fluidity of nucleoli^36^. However, it is important to note that liquid-to-solid transitions are not always undesirable, and indeed may even serve a physiological role^14^. For example, SPD-5, a key protein of the pericentriolar material, forms droplets that become solid-like in minutes^37^, and the resulting material properties mimic those of the pericentriolar material during mitosis^38^. In P granules, MEG-3 forms a gel-like phase^13^.

The gel-like MEG-3 condensates occupy the periphery of P granules, where they were thought to stabilize liquid-like PGL-3 condensates in the middle and help with the localization of P granules in the posterior of *Caenorhabditis elegans* embryos^13^. By contrast, stress granules have stable cores inside a more dynamic shell^8^. Nucleoli have a layered organization where, just as demonstrated using purified proteins, the dense fibrillar component enriched in fibrillarin is embedded inside the granular component enriched in nuleophosmin^10^. In addition, different types of membraneless organelles that share components often physically associate, including Cajal bodies and B snurposomes^39^, P bodies and stress granules^40^, and P granules and P bodies^41^. Physical association has also been observed between droplets formed by unmodified and acetylated chromatin^42^. How the segregation of different condensates in a given membraneless organelle and the physical association of different types of membraneless organelles are maintained is poorly understood.

Evidence suggests that the assembly of membraneless organelles is driven by a few key components^10, 13, 20–30, 33, 37, 43–50^. Other components presumably play regulatory roles, and still others are recruited as clients but may nevertheless have regulatory effects. For the thermodynamics of liquid-liquid phase separation, computational studies led us to the notion that macromolecular regulators fall into three archetypical classes^15, 51^. Volume-exclusion promotors act by taking up space in the bulk phase and thereby displacing driver proteins into the droplet phase. Such promotional effects have been observed in many studies using crowding agents such as Ficoll^20, 23, 25, 27, 31, 33, 37, 48^. Weak-attraction suppressors act by partitioning into the droplet phase and thereby replacing stronger driver-driver attraction with weaker driver-regulator attraction. MEX-5 may have such an effect when helping the posterior localization of P granules^1, 44, 45^. Strong-attraction promotors act in the opposite way, but at high concentrations they turn into suppressors as they overpopulate the droplet phase and repel each other. RNAs are such regulators.

While their promotional effects have been noted in a number of studies^10, 13, 23, 25, 27, 30, 31, 44, 45, 50^, their turnover into suppressors at high concentrations has also been recognized in other studies^21, 34, 46, 49, 52^. The suppressive effects of high-concentrations RNAs may explain their ability to keep ribonucleoproteins soluble^34, 35^.

As support for the forgoing classification of regulators, using pentamers of SH3 domains and proline-rich motifs (SH3_5_ and PRM_5_) as driver proteins^50^, we found Ficoll, lysozyme, and heparin as representatives of volume-exclusion promotors, weak-attraction suppressors, and strong-attraction promotors^51^. This work highlights regulator-driver interaction strength as a key determinant for regulatory effects on thermodynamic properties of phase separation. Macromolecular components can also affect the material properties and time evolution of condensates^19, 32, 37, 42, 47–49, 51^{Maharana, 2018 #75}{Mann, 2019 #21}, but the underlying physical rules are far from clear. Here we uncover some of these rules by studying the phase behaviors of a macromolecular system comprising just four components, SH3_5_, PRM_5_, lysozyme, and heparin.

## Results

Of the four macromolecular components chosen for the present study, SH3_5_ (molecular weight (MW) 43 kD) is a protein composed of five structured domains (i.e., SH3 domains) connected by flexible linkers; PRM_5_ (MW 13 kD) is largely disordered, containing five copies of a proline-rich motif that binds to a specific site on the SH3 domains; lysozyme (MW 14 kD) is a single-domain protein; and heparin (nominal MW 18 kD) is a linear polysaccharide. SH3_5_ has a high negative charge (-28*e*); PRM_5_ has a high positive charge (+30*e*); heparin is extremely anionic, with close to -2*e* per sugar group; and lysozyme has a large positive charge (+8*e*). For notational convenience, we will denote the four macromolecular components as S, P, H, and L, respectively. By mixing different macromolecular components, one can obtain six binaries (e.g., S:P), four ternaries, and one quaternary. As shown below, net charge and structural compactness have significant bearing on the interactions of S, P, H, and L and the phase behaviors of their various mixtures.

### Binary mixtures exhibit three distinct phases

As shown previously^50, 51^, the S:P binary readily forms droplets (Fig. 1a, left). Here we found that S:L and P:H binaries also readily phase separate (Fig. 1a, middle and right; Supplementary Movie 1). In contrast, H:L mixtures form network-like precipitates (Fig. 1b). At a given L concentration (150 μM), the amount of precipitates and, correspondingly, the extent of connectedness of the precipitate networks vary with the H concentration, peaking at 40 μM of H. Whereas S:L and P:H droplets, as observed previously for S:P droplets^51^, fall (under gravity) to the bottom of the solution and then fuse and spread over the supporting coverslip (Fig. 1c and Supplementary Fig. 1a), H:L precipitates initially also fall but then are stable during hours of observation time (Supplementary Movie 2), giving the first hint that H:L interactions are stronger than S:P, S:L, and P:H interactions.

**Figure 1.**
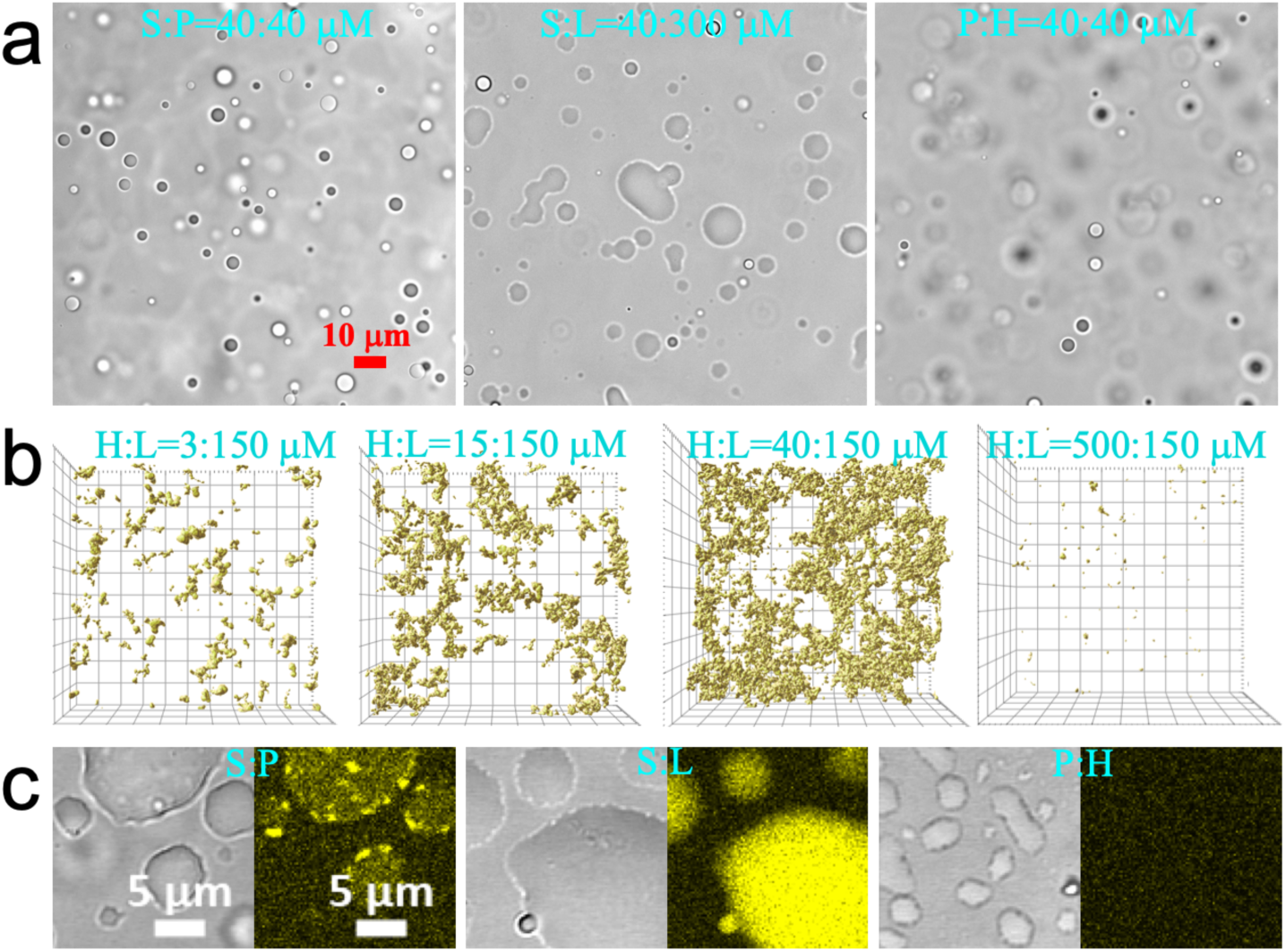
Phases of binary mixtures. (**a**) Brightfield images of droplets formed by S:P, S:L, and H:P binaries, at concentrations indicated. The image of S:P droplets has appeared previously^51^. (**b**) Z stacks of confocal images of H:L precipitates taken at approximately 20 min after mixing H at four concentrations with L at 150 μM, visualized by the fluorescence of thioflavin T (ThT) in a top view. The field of view is 105 μm × 105 μm and the height of the Z stacks range from 30 to 50 μm. (**c**) Brightfield and fluorescence images of S:P, S:L, and H:P droplets, showing moderate, strong, and no ThT binding (yellow), respectively. Concentrations were S:P = 40:40 μM, S:L = 20:300 μM, and P:H = 10:40 μM. Images were taken at 10 to 20 min after mixing; by then droplets had fallen, fused, and spread over a coverslip.

The two remaining binaries, S:H and P:L, stay as a homogeneous solution, consistent with the expectation that two macromolecules with large like charges repel each other and the repulsion prevents condensation. This observation also gives support to the notion that the condensation of the other four binaries, all between oppositely charged macromolecules, is driven to a large extent by electrostatic attraction.

We used thioflavin T (ThT) binding to provide a probe into the binary condensates. ThT emits strong fluorescence when mixed into H:L precipitates and S:L droplets, weak fluorescence when mixed into S:P droplets, and no more than background fluorescence when mixed into P:H droplets (Fig. 1b, c and Supplementary Fig. S1b). As a control, ThT emits only weak fluorescence in an L solution even at 4 mM (Supplementary Fig. S1c). These observations suggest that ThT emits strong fluorescence when bound to structured protein domains that are closely packed, and that the H:L and S:L condensates have very high packing densities, whereas S:P and P:H condensates have intermediate and low packing densities, respectively.

We went on to determine the phase diagrams of the four condensate-forming binaries (Fig. 2). After mixing two types of macromolecules at specified concentrations, observation under a microscope indicated whether condensation occurred, and if so, whether the condensates were droplets or precipitates. The phase boundary for each of the binary is shaped as a tilted parabola, as typified by the S:P phase diagram (Fig. 2b), indicating the need for a proper concentration balance between the two components in condensation. To illustrate, consider L at 150 μM (vertical dashed line in Fig. 2e) for the H:L binary. When the H concentration is ≤ 0.2 μM or ≥ 1500 μM, the H:L binary remains a homogeneous solution. As one moves from the boundaries (H at 0.3 and 1000 μM) toward the middle of the precipitation region, the amount of precipitates increases (Fig. 1b).

**Figure 2.**
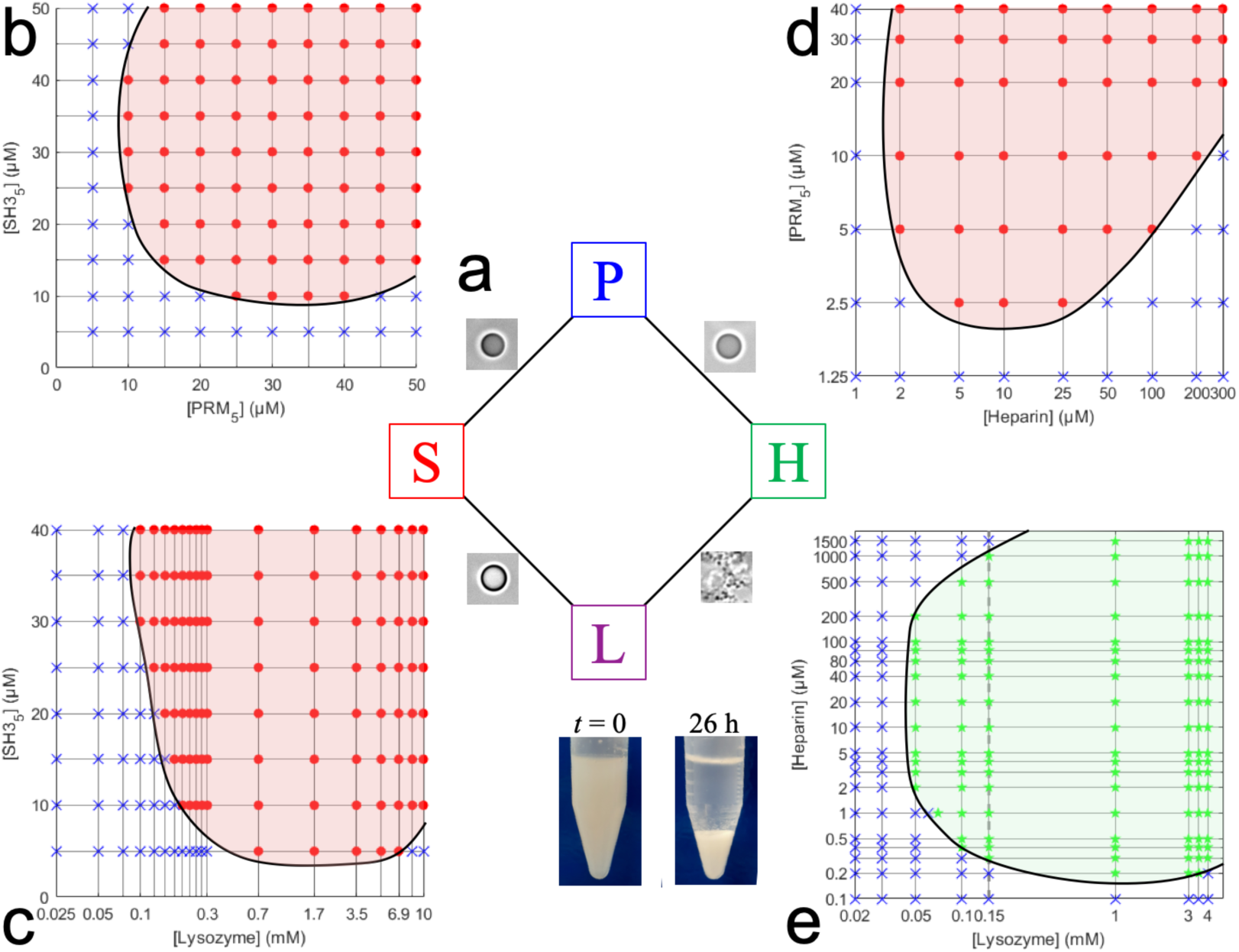
Phase diagrams of four binaries. (**a**) Summary of the condensate phases of the four binaries. (**b-e**) Phase diagrams of S:P, S:L, P:H, and H:L binaries. Red circles, green stars, and blue crosses indicate concentrations at which mixtures form droplets or precipitates, or is a homogeneous solution. Black curves indicate the boundaries between regions where condensation either occurs or does not occur. A vertical dashed line in (**e**) indicates that, at a given L concentration, H concentrations either too high or too low do not result in precipitation. Inset: a solution of 40 μM H and 150 μM L immediately after mixing (*t* = 0) and after 26 h sedimentation under gravity.

Also worth noting are the wider ranges of H and L concentrations leading to H:L precipitation than those leading to P:H and S:L phase separation. In particular, the lowest H concentration for P:H phase separation is 2 μM (Fig. 2d), but the counterpart for H:L precipitation is 10-fold lower, at 0.2 μM (Fig. 2e). Likewise the lowest L concentration for S:L phase separation is 100 μM (Fig. 2c), but the counterpart for H:L precipitation is 2-fold lower, at 50 μM (Fig. 2e). The wider concentration ranges for H:L condensation are another hint that H:L interactions are stronger than P:H and S:L interactions. Still, H:L precipitates can be dissolved by dilution with buffer (into the homogeneous-solution region); this dissolvability distinguishes H:L precipitates from other irreversible aggregates. Moreover, H:L precipitates can be dissolved by increasing the salt (KCl) concentration in the buffer from 150 mM to 500 mM, supporting both the reversibility of the precipitation and the electrostatic nature of the driving force.

Gravity provides a natural way to sediment H:L precipitates. A mixture of 40 μM H and 150 μM L is in the precipitation region (Fig. 2e) and correspondingly appears cloudy immediately after preparation (Fig. 2e inset, *t* = 0). After settling under gravity for 26 h, the precipitates sediment to become a dense, white pellet separated from a clear supernatant (Fig. 2e inset, *t* = 26 h). The L concentration in the supernatant is 46 μM, consistent with the phase boundary determined above; the L concentration in the pellet is 10-fold higher.

### P, but not S, can dissolve H:L precipitates into droplets

We now consider the four ternary mixtures. As found previously^51^, low-concentration H promotes S:P phase separation whereas L always suppresses the phase separation. In doing so, H and L partition significantly into the condensed phase, but the droplets stay as droplets (Supplementary Fig. 2a, b). Similar to the H:L binary, the S:H:L ternary also forms and stays as precipitates (Fig. 3a). Fluorescence images of labeled components show that S and L are distributed throughout precipitate networks but H seems to be more localized (Fig. 3b). ThT also binds uniformly to the precipitate networks (Fig. 3c).

**Figure 3.**
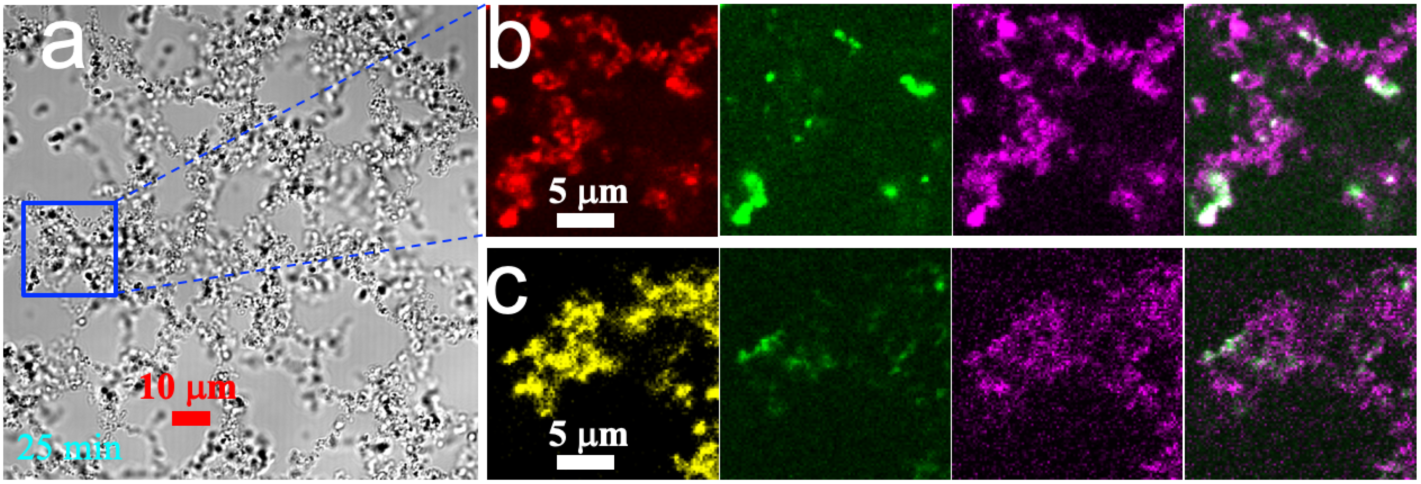
Precipitates formed by an S:H:L ternary mixture. (**a**) Brightfield image of the precipitates at 25 minutes after mixing S, H, and L at 10, 40, and 150 μM, respectively. (**b**) Enlarged view of a region inside the precipitates, visualized by fluorescence of Alexa 594-SH3_5_ (red), FITC-heparin (green), and Cy5-lysozyme (magenta). The last panel shows the merge of the green and magenta channels. (**c**) ThT binding (yellow) to S:H:L precipitates formed from S:H:L = 10:40:150 μM and labeled with FITC-heparin and Cy5-lysozyme, imaged at 5 min after mixing. The last panel shows the merge of the green and magenta channels.

In contrast, the P:H:L ternary can initially form precipitates (Fig. 4a, *t* = 0 min), but over time these precipitates become droplet-like (Fig. 4a, *t* = 10 and 25 min). Indeed, brightfield images of P:H:L condensates at the later time look similar to binary droplets that have fused and spread over a coverslip (compare Fig. 4a, *t* = 25 min and Supplementary Fig. 1a). However, fluorescence images show that the components are not uniformly distributed (Fig. 4b). H and L do not colocalize, although over time they seem to mix. Sometimes, small foci rich in H appear inside a large droplet populated with L (Fig. 4c). ThT binds to the L-populated surround but not the H-rich foci, suggesting that the foci are P:H-rich (see Fig. 1c).

**Figure 4.**
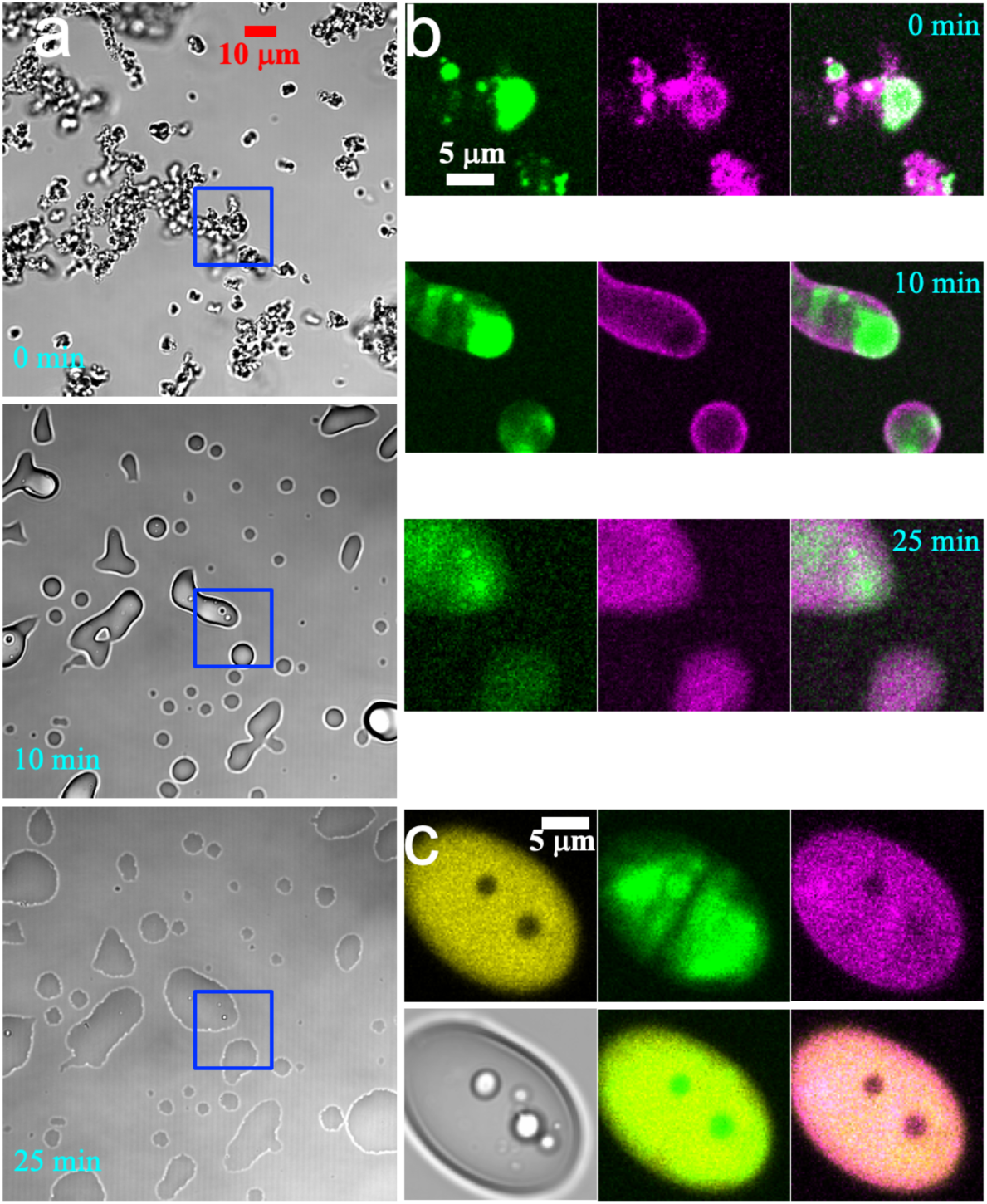
Evolution of P:H:L condensates over time. (**a**) Brightfield images of a P:H:L mixture (10:40:150 μM) at *t* = 0, 10 and 25 min. The region in a blue box is enlarged and shown in (**b**) by fluorescence of FITC-heparin (green) and Cy5-lysozyme (magenta). The merge of the green and magenta channels clearly shows demixing of H and L in the condensates at *t* = 0 and 10 min: FITC-heparin is concentrated in places where Cy5-lysozyme is deficient. At *t* = 25 min, mixing of H and L occurs but is still incomplete. (**c**) ThT binding to a P:H:L condensate at *t* = 5 min after mixing (10:40:150 μM). Top row: images recorded in yellow (ThT), green (FITC-heparin), and magenta (Cy5-lysozyme) channels. Bottom row: brightfield image and merges of the yellow channel with either the green or magenta channel. Within the condensate are two droplets enriched in FITC-heparin but deficient in Cy5-lysozyme and showing poor ThT binding.

### Ternary phase diagrams unveil relative strengths of pairwise interactions

We explored the phase diagrams of the four ternaries. To simplify, when varying the concentrations of the three components, we fixed the ratio between two of the components. The set of three such phase diagrams for the S:P:H ternary is shown in Fig. 5a. The concentration space is divided into one region where droplet formation is observed and another region where the S:P:H ternary remains as a homogeneous solution. In our previous study^51^, we determined H as a strong-attraction promotor of S:P phase separation. Correspondingly, at S:P equimolar ratio, increasing H initially leads to a decrease in the S and P threshold concentration for phase separation but, upon a further increase in concentration, H switches from a promotor to a suppressor and hence the S and P threshold concentration starts to increase (Fig. 5a, left panel). The overall shape of the phase boundary at a fixed S:P ratio is an upward parabola.

**Figure 5.**
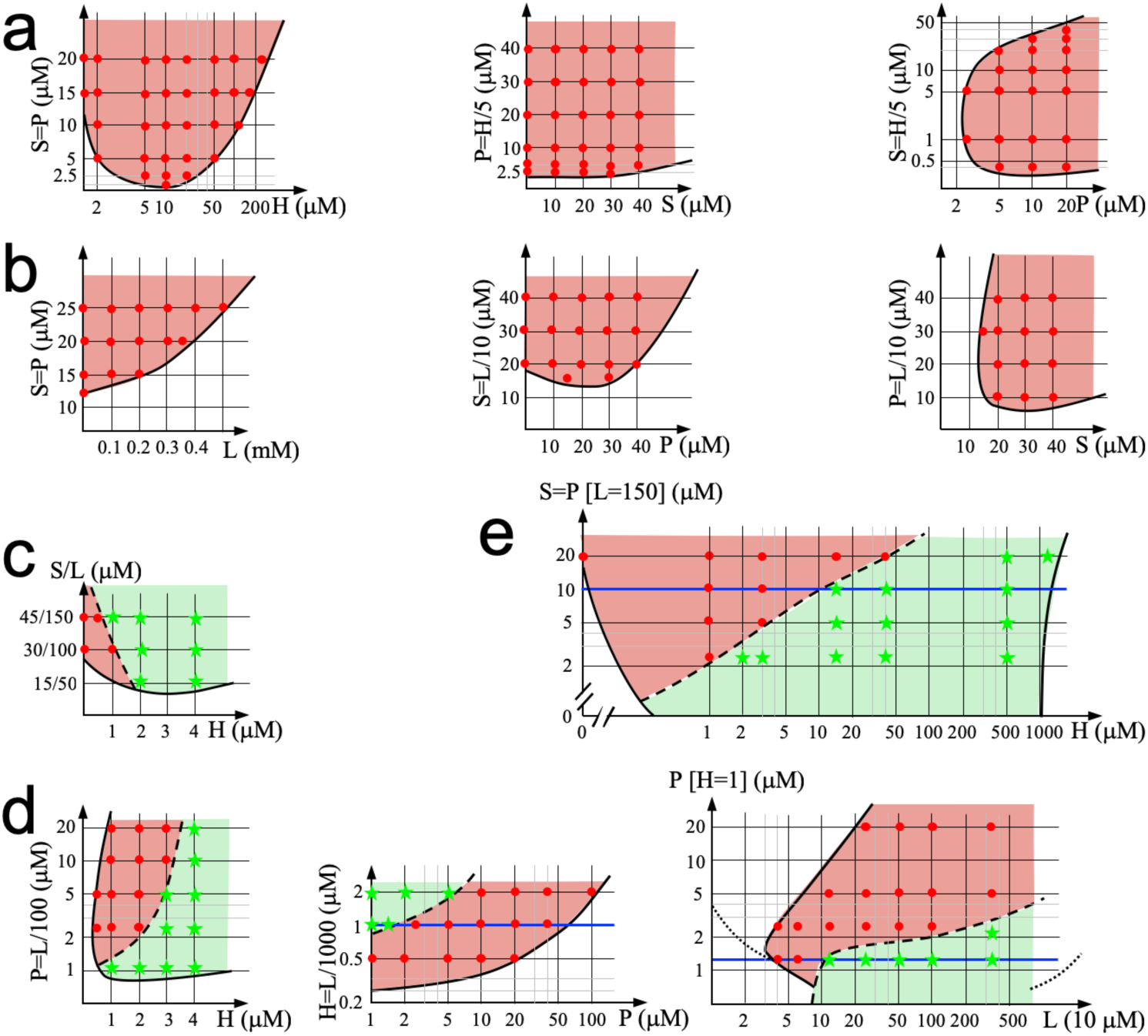
Phase diagrams of four ternaries and the quaternary. (**a**) S:P:H ternary phase diagrams, with the concentration ratio between any two of the three species fixed. The one with the S:P ratio fixed at 1 was amended from a version published previously^51^; the lowest S and P concentrations for droplet formation in the presence of H is now reduced to 1.25 μM. (**b**) S:P:L ternary phase diagrams. (**c**) S:H:L ternary phase diagram, with the S:L concentration ratio fixed at 0.3. The condensate phases are either droplets (light red shading) or precipitates (light green shading), but are predominantly the latter. (**d**) P:H:L ternary phase diagrams, with P:L or H:L ratio fixed or H concentration fixed. In the last phase diagram, the phase boundary would be shaped as an upward parabola (dotted curve) had we fixed the P:H ratio instead of the H concentration. (**e**) Phase diagram of the S:P:H:L quaternary, with S:P ratio and L concentration fixed. A blue horizontal line at S:P:L = 10:10:150 μM indicates that, with H concentration increasing from 0 to beyond 1000 μM, one encounters first solution, then droplet formation and precipitation, and finally back to solution.

Being a strong-attraction promotor implies that the P:H attraction is stronger than the S:P attraction^51^. A corollary of this assertion is that S should be a weak-attraction suppressor of P:H phase separation. This prediction is confirmed by the phase diagram at a fixed P:H ratio (Fig. 5a, middle panel), showing that the threshold concentration for P:H phase separation increases with increasing S concentration. The resulting phase boundary is shaped as a half parabola.

When the ratio of the two negatively charged components, S and H, is fixed, the phase boundary of the S:P:H ternary is shaped as a tilted parabola (Fig. 5a, right panel), similar to the phase boundaries of the binaries (Fig. 2). As explained above, the tilted parabola shape results from the need for a proper concentration balance in condensation, here between the two negatively charged components, S and H, and the positively charged component, P. Considering the three panels of Fig. 5a together, a rule emerges. A ternary of charged species necessarily consists of two components, L1 and L2, with like charges and a third component, O, with an opposite charge. Suppose that the L1:O attraction is stronger than the L2:O attraction. Then, depending on the ratio of which two of the three components is held fixed, the phase boundary is shaped as either an upward parabola, or a half parabola, or a tilted parabola. Specifically, the three shapes are observed when fixing the ratios of the weaker-attraction pair (L2:O), the stronger-attraction pair (L1:O), and the like-charge pair (L1:L2), respectively.

The foregoing ternary rule is immediately confirmed on the S:P:L ternary. Previously we have determined L as a weak-attraction suppressor of S:P phase separation, indicating that S:P attraction is stronger than the S:L attraction. We thus expect the shapes of phase boundaries to be a half parabola, upward parabola, and titled parabola, respectively, when fixing the S:P ratio, the S:L ratio, and the P:L ratio. These are exactly the shapes observed (Fig. 5b).

We can now rank the strengths of the four types of pairwise interactions. Combining the outcomes of the ternary rule on the S:P:L and S:P:H mixtures, we have the following order for the pairwise interactions: P:H > S:P > S:L. The precipitation of the H:L binary has indicated that this pair has the strongest attraction, which is confirmed by the ternary-rule results below for the S:H:L and P:H:L mixtures. The order of the four types of pairwise interactions is thus H:L > P:H > S:P > S:L. The stronger attraction between P and H than between S and L explains why P but not S can dissolve H:L precipitates into droplets.

### Compositional change produces a tug of war between phases

The S:H:L ternary presents an interesting case because it may form droplets when S and L are in excess over H but precipitates when H and L are in excess over S. Therefore the concentration space is divided into three regions: one where droplet formation is observed; another where precipitation is observed; and a third where a homogeneous solution is observed. However, for S:H:L, the droplet-forming region is very small and condensation is dominated by precipitation (Fig. 5c). The dominance yet again implicates much stronger H:L attraction than S:L attraction. The stronger H:L attraction is further confirmed by the upward parabolic shape of the boundary for the combined regions of condensation (including both droplet formation and precipitation), according to the ternary rule.

The P:H:L ternary proves to be a more even competition between droplet formation and precipitation (Fig. 5d). When the P:L ratio is fixed, the boundary for the combined regions of condensation is shaped as a tilted parabola (Fig. 5d, left panel), as expected from the ternary rule. The droplet and precipitation regions are divided approximately equally. Likewise, when holding the H:L ratio constant, the phase boundary for condensation has a half-parabola shape (Fig. 5d, middle panel), consistent with the ternary rule and confirming stronger H:L attraction than P:H attraction. Along a horizontal line (blue line in Fig. 5d, middle panel), precipitates occur at low P whereas droplets occur at high P. These data plainly indicate that increasing amounts of P dissolve precipitates into droplets. This finding is reinforced by the approximately 45° slope of the dividing line between the droplet and precipitation regions for the P:H:L ternary (Fig. 5d, left and middle panels), in contrast to a 145° slope for the S:H:L ternary (Fig. 5c); the 45° slope means dissolution of precipitates by increasing amounts of P even as H and L concentrations are also increasing. The competition between precipitation and droplet formation thus can occur both over time (Fig. 4) and with a change in macromolecular composition. The classification of P:H:L condensates into precipitates is based on the observation at *t* = 0 (i.e., right after the components are mixed). All these precipitates, similar to those shown in Fig. 4, have the tendency to become droplet-like over time.

We also determined the phase diagram of the P:H:L ternary while holding H at 1 μM (Fig. 5d, right panel). At this H concentration, the P:H binary does not phase separate no matter the P concentration, but the H:L binary precipitates for L above 0.7 μM (Fig. 2). Points along a horizontal line (blue line in Fig. 5d, right panel) illustrate the tug of war between phases. With P held at 1.25 μM, the P:H:L ternary starts to form droplets when L is at 40 μM but changes into precipitates when L is at 125 μM; finally the P:H:L ternary is expected to be a homogeneous solution at very high L (> 5 mM), similar to a pure L solution. As the L concentration increases, the winner in the tug of war changes hands multiple times, first from homogeneous solution to droplets, then from droplets to precipitates, and finally from precipitates back to homogeneous solution.

The tug of war between phases is also observed in the S:P:H:L quaternary (Fig. 5e). While holding L at 150 μM and the S:P ratio at 1, we observe droplet formation when S and P are in excess over H whereas precipitation when H is in excess over S and P. Along a horizontal line (blue line in Fig. 5e), we again see the winner changing hands from homogeneous solution to droplets at low H, then from droplets to precipitates at intermediate H, and finally from precipitates back to homogeneous solution at high H. Along a vertical line, the points show that increasing amounts of S and P together dissolve H:L precipitates into droplets.

### Mixing of S:P droplets with H:L precipitates directly demonstrates dissolution

The last observation suggests that H:L precipitates can be dissolved when they encounter S and P. To test this idea, we placed a 1 μl drop of H:L precipitating solution (concentrations at 40:150 μM and labeled with FITC-lysozyme) in contact with a 1 μl drop of S:P droplet-forming solution (concentrations at 25:25 μM and labeled with Alexa 594-SH3_5_). As Supplementary Movie 3 shows, H:L precipitates become droplet-like, either quickly when they fuse with S:P droplets, or a bit more slowly when they take up S and P from the bulk solution. This timelapse video is composed of slices, with decreasing *Z* positions, each selected from a *Z* stack in a time series, and hence is called *Z*-chase.

### S:P:H:L precipitates become droplet-like over time but display demixing of components

When S and P are below the concentrations for dissolving H:L precipitates, the S:P:H:L quaternary forms precipitates, which, like the P:H:L counterparts (Fig. 4) evolve over time in material state (Supplementary Movies 4 and 5). However, the situation with the quaternary system is more complex and interesting (Fig. 6).

**Figure 6.**
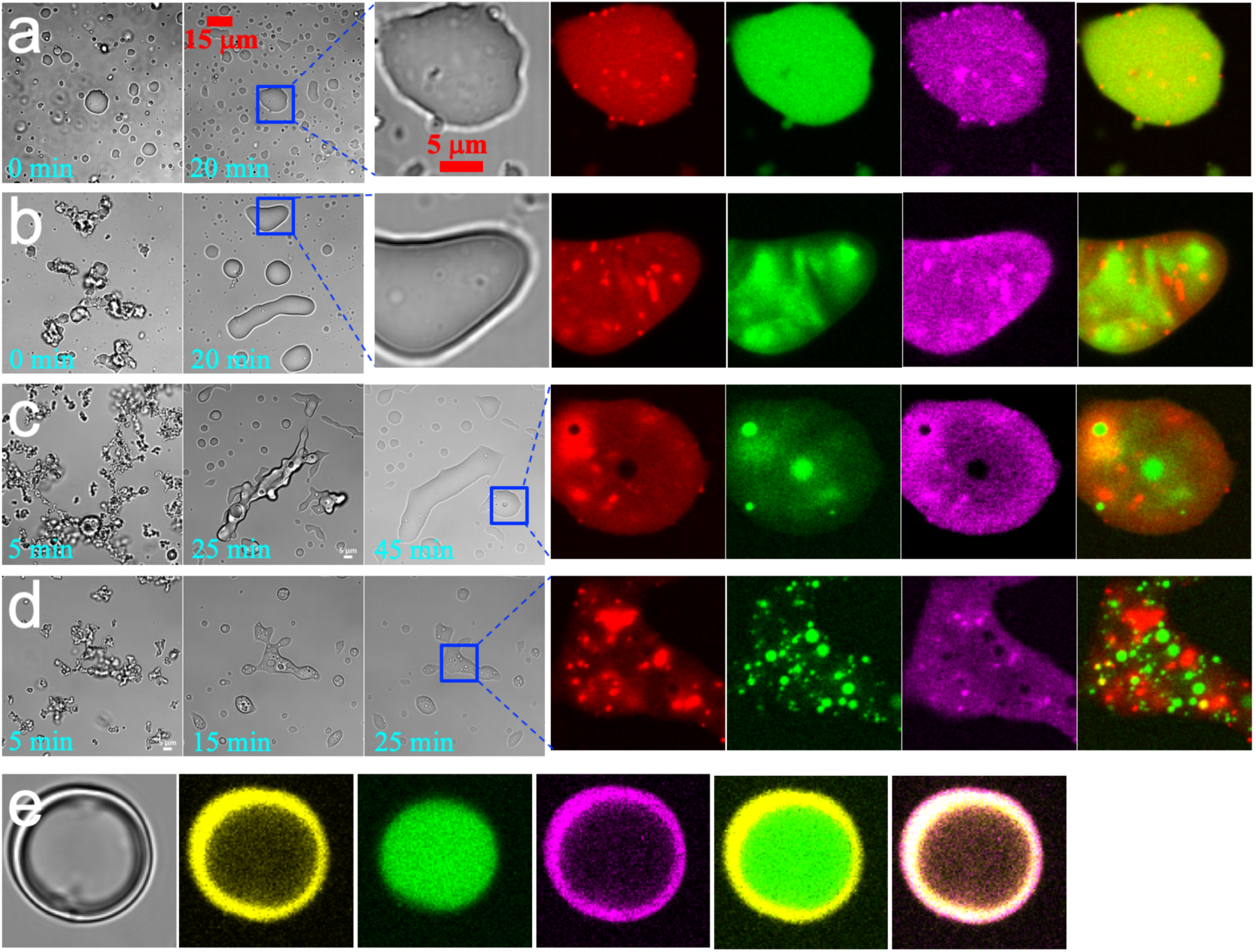
Time evolution of and demixing in S:P:H:L condensates, at S:P:L = 10:10:150 μM. (**a-d**) Brightfield and fluorescence images of the quaternary with H = 3, 15, 40, and 500 μM at indicated times. The samples were labeled with Alexa 594-SH3_5_ (red), FITC-heparin (green), and Cy5-lysozyme (magenta). (**e**) ThT binding to a condensate (S:P:H:L = 10:10:500:150 μM) at *t* = 5 min. From left to right: brightfield image, images recorded in yellow (ThT), green (FITC-heparin), and magenta (Cy5-lysozyme) channels, and merges of the yellow channel with either the green or magenta channel.

To illustrate the complex phase behaviors of the quaternary system, we observed samples on a confocal microscope at four points along the blue line in Fig. 5e, with S:P:L = 10:10:150 μM but increasing H. These samples were labeled with Alexa 594-SH3_5_ (red) and FITC-lysozyme (green). At the first point, H = 3 μM and the quaternary forms droplets, which like S:P^51^ and other binary droplets, fall under gravity, and then fuse and spread over a coverslip (Supplementary Movie 4a). The other three points, at H = 15, 40, and 500 μM, are in the precipitating region. Once precipitates are formed, while falling they fuse into condensates with a smooth surface (Supplementary Movies 4b-d). Initially the precipitate networks look similar to those formed with H and L alone (Fig. 1b), except that the amount of precipitates at H = 500 μM is noticeably larger in the presence of S and P, indicating that the latter components expand the region of condensation (see this expansion at increasing S and P concentrations in Fig. 5e). The most important difference though is that, whereas the H:L precipitates are stable (Supplementary Movie 2), the S:P:H:L precipitates become droplet-like over time. The precipitates at H = 40 μM are the highest in total amount among the three H concentrations, and correspondingly take the longest to transform into droplet-like condensates. The brightfield version of a *Z*-chase video (Supplementary Movie 5), produced from the same time series of Z stacks played in Supplementary Movie 4c, captures this dramatic transformation from a solid-like state to a liquid-like state. However, the fluorescence version, using the merge of the Alexa 594-SH3_5_ (red) and FITC-lysozyme (green) channels, shows red foci inside the condensates, emerging at the very beginning and remaining throughout the transformation in material state, clearly indicating demixing of components.

To more closely inspect the demixing of components, we prepared S:P:H:L samples labeled with Alexa 594-SH3_5_ (red), FITC-heparin (green), and Cy5-lysozyme (magenta). Brightfield images of these samples at different time points show the same evolution in material state as just described (Fig. 6a-d). Even in these brightfield images, many small droplets, or foci, are visible inside large droplets or droplet-like condensates. Fluorescence images further show the main components of the foci. In particular, at H = 3 μM, S (red) and L (magenta) colocalize in tiny foci inside a large droplet, but H (green) distributes uniformly within the large droplet. At H = 15 μM, in addition to S:L-rich foci, there are regions rich in H but poor in S and L. At H = 40 μM, there are now distinct S:L-rich foci and H-rich foci; the latter in all likelihood are also rich in P (see next paragraph). Finally at H = 500 μM, distinct S:L and P:H foci become even more populated. Overall, with increasing H, there is an increasing tendency for S to preferentially mix with L and H to preferentially mix with P, and the two types of binary mixtures segregate, as if undergoing a second phase separation, inside the droplet-like condensates.

As a further probe of the component demixing, we studied ThT binding in these condensates, formed at S:P:H:L = 10:10:500:150 μM with FITC-heparin and Cy5-lysozyme labels (Fig. 6e). An H-rich droplet inside a condensate shows poor ThT binding, whereas the surrounding L-rich region shows strong ThT binding. These observations, considered in light of the contrasting behaviors of S:L and P:H droplets in ThT binding (Fig. 1c), provide strong support to the suggestion that the L-rich regions are mixed with S whereas the H-rich regions are mixed with P.

One possible reason why the demixing happens is that S:L droplets and P:H droplets do not mix with each other. To check this possibility, we placed a 1 μl drop of S:L droplet-forming solution (concentrations at 30:300 μM and labeled with Cy5-lysozyme) in contact with a 1 μl drop of P:H droplet-forming solution (concentrations at 40:40 μM and labeled with FITC-heparin). To partly mimic the macromolecular contents inside the S:P:H:L condensates, we even added 100 g/L Ficoll70 to these solutions. When an S:L droplet falls on top of a P:H droplet, they start to fuse (Supplementary Fig. 3a). By 5 min the contents are completely mixed (Supplementary Fig. 3b). Meanwhile S:L droplets also take up H (and P as well; see Fig. 4b, middle panel) from the bulk solution, and P:H droplets take up L (and S as well; see Fig. 4a, middle panel) from the bulk solution. Clearly, S:L and P:H droplets, even in a macromolecular background provided by Ficoll70, can quickly exchange contents. Ficoll70 is not “sticky” to these macromolecular components and, as shown previously^51^, does not partition appreciably into S:P droplets.

It thus seems most likely that the macromolecular background inside S:P:H:L condensates forms a matrix that is “sticky” to S:L-rich and P:H-rich mixtures. The stickiness hinders the dissociation of components from foci and their diffusion within the background matrix. The rapid emergence of foci upon mixing of the components may also be facilitated by this stickiness. Supporting this conclusion, when S:P:H:L condensates from two solutions, one labeled with FITC-lysozyme and the other with Cy5-lysozyme, are mixed, fusion is stalled (as evidenced by irregular shapes of fused condensates), and foci initially formed in the two solutions persist and are only diluted slowly by infusion of components from the bulk solution (Supplementary Movie 6).

### FRAP shows hindered diffusion of components in S:P:H:L condensates

Fluorescence recovery after photobleaching (FRAP) provides a direct way to demonstrate hindered diffusion. As reference, fluorescence inside S:P droplets (formed at 40:40 μM) recovers quickly, to approximately 70% of the pre-bleach level (Fig. 7a; Fig. 7b, first row). S:P:H:L droplets (formed at 10:10:3:150 μM) behave in much the same way (Fig. 7a; Fig. 7b, second row). In contrast, fluorescence inside S:P:H:L condensates (Fig. 7c) that started as precipitates (formed when H is elevated to 15, 40, or 500 μM) fail to recover to a significant extent (Fig. 7a; Fig. 7b, third to fifth row; Fig. 7c). Indeed, the extents of recovery in these precipitates-turned condensates are similar to that inside S:H:L precipitates (Fig. 7a; Fig. 7b, last row). Therefore, despite the transformation from a solid-like state to a liquid-like state in outward appearance, the interior of these condensates is not completely liquid as diffusion of components is hindered, a feat befitting to the persistent demixing of components.

**Figure 7.**
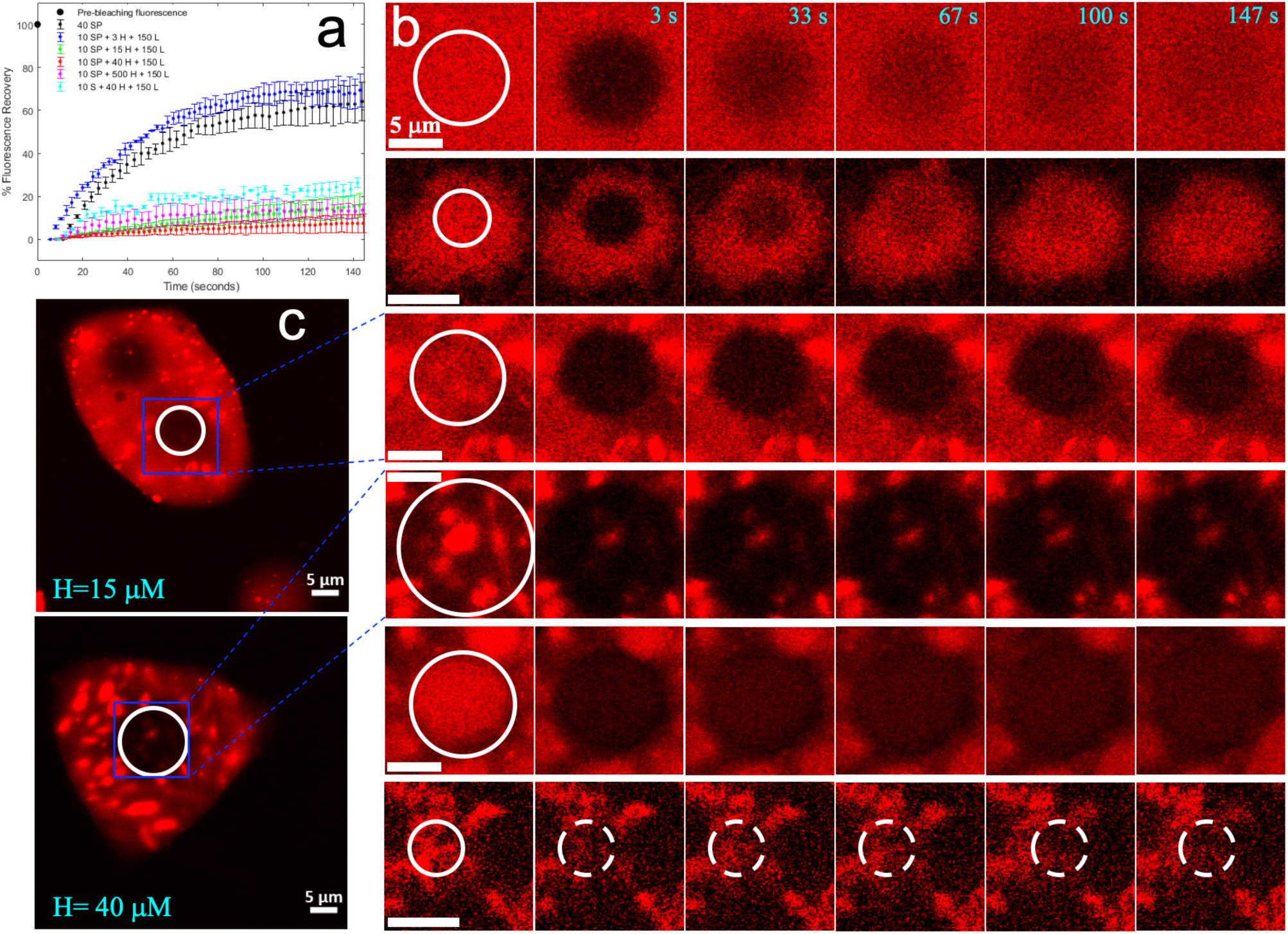
Fluorescence recovery after photobleaching (FRAP) of binary, ternary, and quaternary mixtures. (**a**) Time dependences of the percentages of fluorescence intensity recovered in different condensates. The condensates were formed by mixing species at concentrations (in μM) shown in the legend, and had settled over a copverslip. (**b**) Fluorescence images before bleaching and at indicated times after bleaching. White circles indicate bleached regions. From top to bottom: S:P, S:P:H:L at H = 3, 15, 40, and 500 μM, and S:H:L. (**c**) Post-FRAP images of two S:P:H:L mixtures over a larger field containing the cropped regions shown in (**b**). The image at H = 15 μM was taken 10 min after bleaching of the circled region, still showing no recovery. To the upper left is another spot where FRAP was done even earlier, by 5 min. The image at H = 40 μM was taken immediately after completion of FRAP in the circled region.

## Discussion

Using four macromolecular components alone, we have demonstrated complex phase behaviors associated with membraneless organelles and probed the underlying physical rules. The dissolution of heparin:lysozyme precipitates by SH35 and PRM5 into droplets may serve as a model for macromolecular regulation of the delicate balance between physiological membraneless organelles and aberrant aggregation^20–31^. While different bimolecular condensates have been reported to undergo liquid-like to solid-like aging^14^, either as a pathological process or for normal function, the transformation from solid-like precipitates to droplet-like condensates exhibited by our quaternary system represents a time evolution of material properties in the opposite direction. Our findings may have implications for the disassembly of solid-like biomolecular condensates^32–35^ and pathological aggregates such as Aβ fibrils. Lastly the demixing of SH3_5_:lysozyme-rich and PRM_5_:heparin-rich foci inside the droplet-like condensates may be instructive for understanding the existence of stable cores inside stress granules^8^ and the segregation of different condensates in nucleoli^10^ and P granules^13^.

To deconstruct the complex phase behaviors of the quaternary system, we have studied the binary and ternary mixtures of the components. That the heparin:lysozyme binary forms precipitates but the other three binaries form droplets sets off a tug of war between phases in the ternary and quaternary systems. The contrasting phases of the four binaries can be explained by differences in intermolecular interaction strength and degree of structural compactness, in line with previous studies^17–19^. Importantly, we have found that the strengths of pairwise interactions can be ranked by determining ternary phase diagrams. The ternary rule developed here should have broad applications. According to this rule, the order of the strengths of four pairwise interactions is: heparin:lysozyme > PRM_5_:heparin > SH3_5_:PRM_5_ > SH3_5_:lysozyme. This order helps explain not only why the heparin:lysozyme binary forms precipitates but also why PRM_5_, but not SH3_5_, by itself can dissolve heparin:lysozyme precipitates. The pairwise attractive interactions, mostly electrostatic in nature, are responsible for the stickiness of the macromolecular matrix in quaternary condensates. The differences in the afore-mentioned two factors, interaction strength and structural compactness, as well as the ensuing competition among the macromolecular components also contribute to the transformation from precipitates to droplet-like condensates and the demixing of components in the condensates. The common electrostatic nature for the different material states observed in the present study challenges the view that electrostatic interactions confer liquid-like properties whereas hydrophobic interactions confer solid-like properties^53^.

Our understanding of the quaternary system and its subsystems, based on determination of phase diagrams, confocal microscopy, ThT binding, mixing experiments, and FRAP, can be summarized by a set of cartoons illustrating the molecular interactions and organizations inside the condensates (Fig. 8). For the four types of binary condensates, ThT binding has revealed different packing densities, related to the structural compactness of the components. In an S:P droplet, each S (a string of five loosely connected domains) binds multiple copies of P (an unstructured polymer) and vice versa. The binding is weak, and so the partners constantly unbind and rebind, producing a liquid state. The same description holds for S:L and H:P droplets, except that S wraps the single structured domain of L leading to closer packing, and the unstructured polymers P and H act as physical crosslinks of each other leading to looser packing.

**Figure 8.**
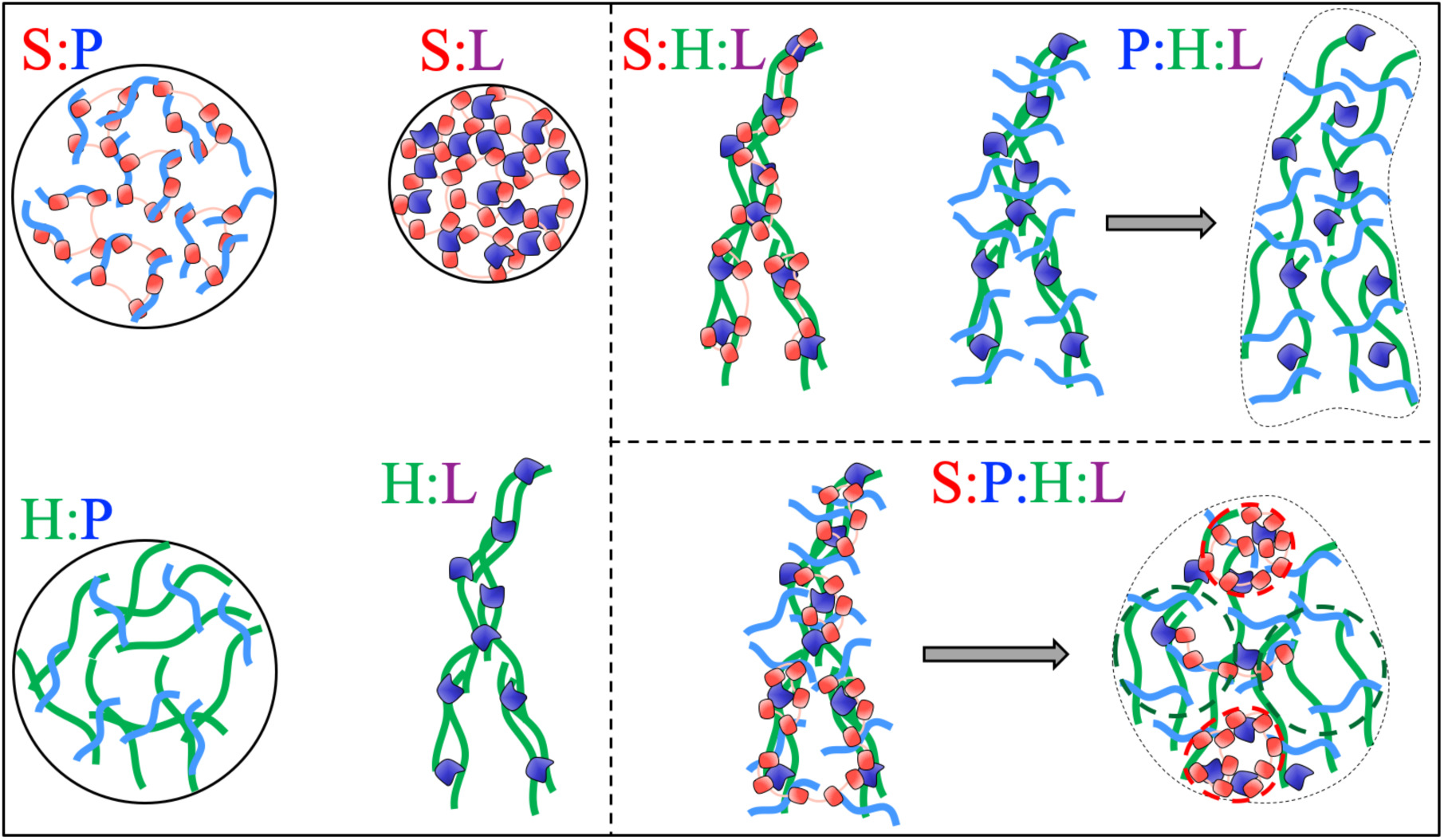
Cartoons illustrating molecular arrangements inside binary, ternary, and quaternary condensates. For P:H:L and S:P:H:L condensates, a horizontal arrow indicates evolution over time.

In H:L precipitates, H polymers are physically crosslinked by multiple copies of the small L domain, leading to tight packing and hence a solid-like state. The S:H:L precipitates are similar, except that now S also wraps around L. The P:H:L ternary initially forms precipitates, in which P binds to parts of H that are not already covered by L. As P and H prefer looser packing, the precipitate networks expand locally and branches of the networks become connected, leading to a liquid-like state. The implication of both interaction strength and degree of structural compactness as determinants of material properties unites the findings from two recent studies^19, 48^. S:P:H:L precipitates undergo a similar transformation from a solid-like state to a liquid-like state; here, in the H:L initiated precipitates, S preferentially binds to L, whereas P preferentially binds to uncovered parts of H, thus generating S:L-rich and P:H-rich foci inside the quaternary condensates.

## Materials and Methods

### Protein expression, purification, and labeling

Expression and purification of SH3_5_ and PRM_5_ were as described previously^51^, and so was the labeling of SH3_5_ (Alexa 594-SH3_5_) with Alexa Fluor 594 NHS ester. Labeling for PRM_5_ was not attempted, because that would modify lysine side chains important for macromolecular interactions.

### Procurement of other macromolecules

Sources for lysozyme, heparin, FITC-lysozyme, FITC-heparin, and Ficoll70 were the same as reported previously^51^. Cy5-lysozyme was obtained as a solution from Nanocs Inc. (catalog #LS1-S5-1), and dialyzed into the final buffer, which contained 10 mM imidazole pH 7, 0.01% NaN_3_, and 150 mM KCl. This buffer was used for all the experiments. Protein concentrations were measured using a NanoDrop spectrophotometer. For heparin, stocks at 5 and 30 g/L were prepared by weighing required amounts on an analytical balance and making up to desired volumes with buffer.

### Determination of phase diagrams

All phase diagrams were constructed by preparing a binary, ternary, or quaternary mixture from the four components, lysozyme, PRM_5_, SH3_5_ and heparin, in a microcentrifuge tube. The components were added, always in that order, as quickly as possible and mixed thoroughly using a pipette (see Supplementary Movie 1). The samples were then immediately observed under an Olympus BX53 microscope using a 40x/0.75 UPlan FL-N objective, and recorded as either homogeneous solution, or droplet formation, or precipitation. In cases where condensates changed material states over time, the results reported are for *t* = 0. The phase diagrams were plotted using MATLAB and the boundaries were drawn manually.

### Confocal microscopy

The protocol for sample preparation was the same as previously described^51^. Images stamped *t* = 0 were captured immediately after the slide was mounted to the microscope. In cases where observations were made over a period longer than an hour, sample evaporation was minimized by sealing the edges of the coverslip with rapid-dry nail-polish topcoat (Electron Microscopy Sciences, catalog #72180).

Brightfield images for the binary systems were acquired on a Zeiss Axio Observer with an alpha Plan-FLUAR 100x/1.45 oil objective. The images were recorded over a 133.12 µm × 133.12 µm field, at a 2048 by 2048 pixel resolution and with a 100 ms exposure time.

All fluorescence images (along with the corresponding images in the brightfield channel), 4D videos, and brightfield and fluorescence videos were taken on a Zeiss LSM 710 with a C-Apochromat 40x/1.2 water objective. The concentrations of the labeled species for fluorescence imaging (unless otherwise indicated) were as follows: FITC-lysozyme and Cy5-lysozyme at 5 µM, FITC-heparin at 3 µM, and Alexa 594-SH3_5_ at 1 µM. The concentration reported for any component was the sum of the unlabeled and labeled (if present) species. The ThT concentration was always 40 µM. For fluorescence images, a Z-stack scanning at a step size of 0.48 µm was conducted, with the first slice set into the coverslip and the last slice at 30 - 50 µm above, enough to cover most of the condensates after brief initial falling under gravity. Each slice covered a 105 µm × 105 µm field with a 512 by 512 pixel resolution, and was scanned four times to obtain average intensities, a number chosen as a compromise between spatial and temporal resolutions.

For ThT fluorescence images, the excitation wavelength was 405 nm and detection wavelength range was 475-600 nm. When ThT-containing samples were also labeled with FITC-heparin and Cy5-lysozyme, track 1 was used for ThT (405 nm excitation and 494-589 nm detection), while track 2 was used for both FITC-heparin (488 nm excitation and 494-589 nm detection) and Cy5-lysozyme (633 nm excitation and 642-727 nm detection). For samples labeled with FITC-heparin (or FITC-lysozyme), Alexa 594-SH3_5_, and Cy5-lysozyme, detections of these species were assigned, respectively, to track 1 (488 nm excitation and 494-589 nm detection), track 2 (561 nm excitation and 592-651 nm detection; unused if Alexa 594-SH3_5_ was absent), and track 3 (633 nm excitation and 665-735 nm detection; unused if Cy5-lysozyme was absent).

### Fluorescence recovery after photobleaching

FRAP experiments were conducted on a Zeiss LSM 710, in triplicates at time points 5 to 10 min apart and in different regions of interest (ROIs), after condensates, labeled with Alexa 594-SH3_5_, had settled on the coverslip. Following two scans, circular ROIs with 3–12 µm diameters were bleached with a 561 nm laser 100 times over 0.9 s at 75% power. The mean intensities in the ROI, obtained in a single recovery scan, were used to calculate percent recovery, and the average of the values from the triplicate measurements was plotted in MATLAB, with standard deviations drawn as error bars.

### Preparation of videos

3D rendering of *Z* stacks from the fluorescence of ThT mixed in H:L samples was done using Imaris 9.5.0 as described previously^51^. *Z* stacks in a time series, each consisting of FITC-lysozyme fluorescence data in S:P:H:L samples, were processed in a similar way, and exported as a set of tiff files, which was then converted into a video in MATLAB.

Brightfield timelapse videos were produced in MATLAB by collating raw images from the transmitted light channel, each for a slice from a *Z* stack in a time series. For an H:L sample, the *Z* positions of the selected slices were fixed. For other samples, the Z positions decreased over time, to account for the falling of condensates under gravity. We refer to such timelapse videos “*Z*-chase”. For the latter samples, a fluorescence version of Z-chase videos was also generated, with images from the merge of FITC-lysozyme (green) and Alexa 594-SH3_5_ (red) channels.

### Mixing experiments

For the mixing of H:L precipitates with S:P droplets, an H:L precipitating solution (concentrations at 40:150 μM and labeled with 10 μM FITC-lysozyme) and an S:P droplet-forming solution (concentrations at 25:25 μM and labeled with 2 μM Alexa 594-SH3_5_) were separately prepared. A 1 μl drop was then pipetted from each solution and placed in contact with each other on a slide. Observation on a Zeiss LSM 710 was as described above, except for the following details: (1) the interface between the two drops was placed near the center of the field of view; (2) the number of scans of each slice was 2 instead of 4; and (3) the detection wavelength range for Alexa 594-SH3_5_ was 602-700 nm.

A similar protocol was followed for mixing S:L droplets with P:H droplets. Concentrations of the initial S:L and P:H solutions were 30:300 μM (labeled with 10 µM Cy5-lysozyme) and 40:40 μM (labeled with 6 µM FITC-heparin), respectively. These solutions each also contained 100 g/L Ficoll70, to slow down diffusion. The detection wavelength range for Cy5-lysozyme was 642-727 nm.

For the mixing of differently labeled S:P:H:L condensates, two solutions were prepared at S:P:H:L = 10:10:40:150 μM, one containing 10 μM FITC-lysozyme and the other 10 μM Cy5-lysozyme. A 1 μl drop was pipetted from each solution and placed on top of each other over a slide. The field was scouted to find regions where both FITC-lysozyme- and Cy5-lysozyme-labeled condensates were visible. From 7 minutes after the slide was mounted, a time series of *Z* stacks was recorded. Each stack consisted of 32 slices, separated by 1 μm in *Z* positions. The detection wavelength range for FITC-lysozyme was 495-581 nm and that for Cy5-lysozyme was 642-735 nm.

## Acknowledgments

This work was supported by National Institutes of Health Grant GM118091.

## Author contributions

H.X.Z. and A.G. designed the research. A.G. and X.Z. performed the research and analyzed the data. H.X.Z. and A.G. wrote the manuscript.

