## Supplementary Figures for "A Tug of War Between Condensate Phases in a Minimal Macromolecular System"

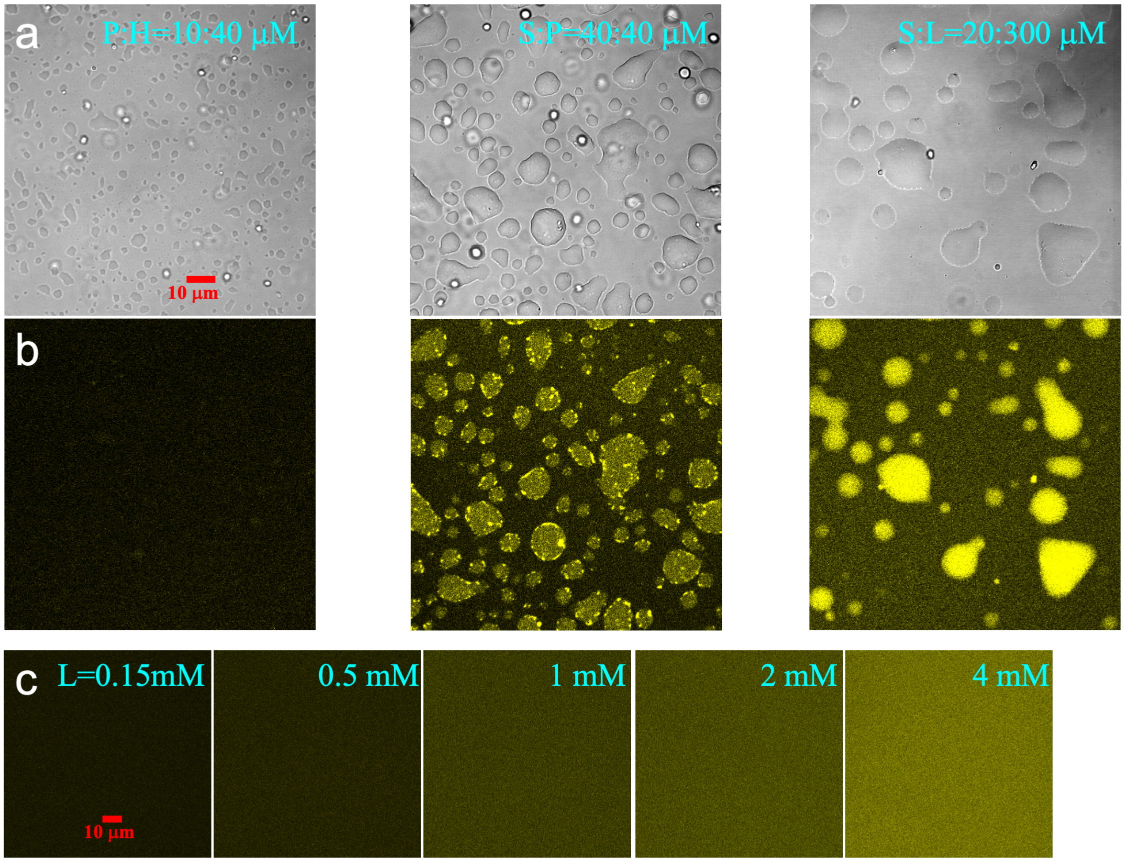


**Supplementary Figure 1 | Different extents of ThT binding to P:H, S:P, and S:L droplets and to L in solutions at varying concentrations.** (**a**) Brightfield images of P:H, S:P, and S:L droplets after fusion and spread over a coverslip. (**b**) Images of the corresponding droplets shown by ThT fluorescence, indicating no, moderate, and strong ThT binding. Images shown here were taken over a large field of view, from which a cropped region is shown in Fig. 1c. (**c**) Fluorescence images of ThT mixed into five L solutions at concentrations ranging from 0.15 to 4 mM.

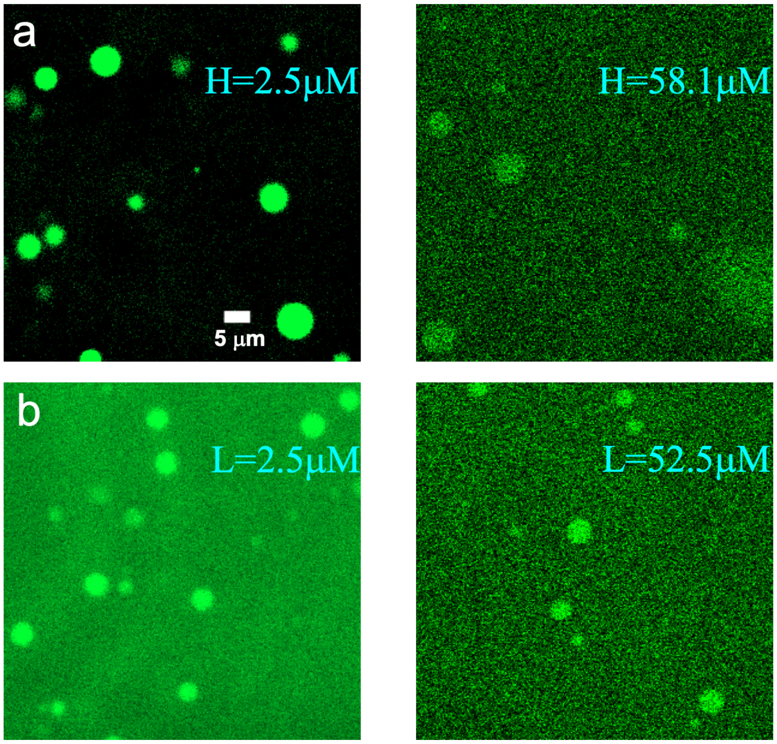


**Supplementary Figure 2 | Images of S:P:H and S:P:L droplets, shown by fluorescence of FITC-heparin and FITC-lysozyme, with S:P = 40:40 μM.** (**a**) S:P:H, with H at 2.5 or 58.1 μM. (**b**) S:P:L, with L at 2.5 or 52.5 μM. Here the concentration of each labeled species was 2.5 μM; images at H = 2.5 μM and at L = 2.5 μM have appeared previously [Ghosh et al. Proc Natl Acad Sci USA 116, 194740-19483 (2019)].

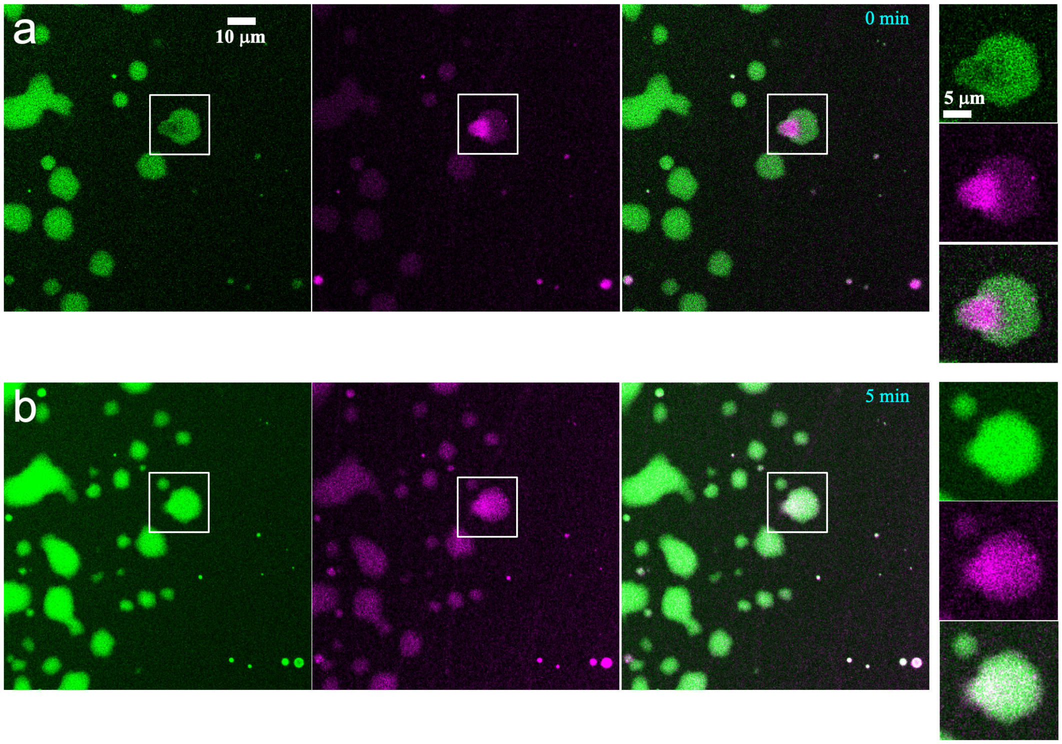


**Supplementary Figure 3 | Images of P:H droplets mixing with S:L droplets, shown by fluorescence of FITC-heparin (green) and Cy5-lysozyme (magenta).** (**a**) After placing two drops, containing P:H and S:L droplets, respectively, side by side (the former is on the left and the latter on the right), a P:H droplet falls on top of an S:L droplet (region inside white box). From left to right: green channel, magenta channel, their merge, and enlarged views of the boxed region. (**b**) Corresponding images after 5 min, showing fusion and content mixing of the P:H and S:L droplets. P:H = 40:40 μM and S:L = 30:300 μM. The drops also contained 100 g/L Ficoll70.
